# Evaluation of a set of eggplant (*Solanum melongena*) lines with introgressions of *S. incanum* under water stress conditions

**DOI:** 10.1101/2023.05.04.539376

**Authors:** Martín Flores-Saavedra, Pietro Gramazio, Santiago Vilanova, Diana M. Mircea, Mario X. Ruiz-González, Óscar Vicente, Jaime Prohens, Mariola Plazas

## Abstract

As access to irrigation water becomes increasingly limited, introgression of relevant genomic regions from drought-tolerant wild genotypes is a promising breeding strategy for crop plants. In this study, nine eggplant (*Solanum melongena*) introgression lines (ILs) covering altogether 71.6% of the genome of the donor wild relative parent *S. incanum* were evaluated for drought tolerance under water stress conditions. Plants at the five true leaves stage were irrigated at either 100% (control) or 30% (water stress) field capacity for 14 days, and growth and biochemical traits were measured. Reduced irrigation resulted in decreased growth and increased levels of stress markers such as proline and malondialdehyde. Most ILs had lower growth and biomass production than the cultivated parent under both conditions. However, the wild alleles for two QTLs related to stem and root dry weight (*dwt8* and *dwr6%*) conferred improved tolerance to water stress. In addition, several *S. incanum* alleles had a positive effect on important traits that may improve yield under drought conditions, such as leaf water content (*lwc12%*), water use efficiency (*wue1%*) and chlorophyll content (*chl2* and *chl8%*). Fine-mapping of the QTLs for tolerance and reducing linkage drag with regions affecting growth will be crucial for significantly improving eggplant drought tolerance through introgression breeding.

## 1. Introduction

The effects of climate change, including reduced rainfall and increased drought, are expected to pose significant challenges to agriculture in many world regions (Arnell et al., 2019). The development and use of drought-tolerant genotypes will be crucial for adapting to these dramatic changes. Among vegetable crops, eggplant (*Solanum melongena* L.) is particularly important in tropical and subtropical regions, which are expected to be the most affected by climate change (Chapman, 2020). Although eggplant is considered tolerant to mild water stress and can thrive with reduced irrigation levels without significant impact on yield (Díaz-Pérez & Eaton, 2015), some of its wild relatives growing in arid and semi-arid regions have shown higher tolerance to oxidative stress and more promising results under limited water conditions (Plazas et al., 2022).

The use of introgression lines (ILs) is a powerful tool for genetic improvement and quantitative trait locus (QTL) identification, allowing the evaluation of the potential of individual introgressions from wild species in an otherwise cultivated genetic background (Lippman et al., 2007). The development and use of ILs with wild introgressions have allowed the improvement of tolerance to abiotic stress conditions in several crops, such as rice (Ali et al., 2017) or pearl millet (Sharma et al., 2020). In tomato, drought tolerance was improved using rootstocks derived from ILs of the wild species *S. habrochaites* S. Knapp & D.M. Spooner (Poudyal et al., 2017).

In eggplant, a set of ILs developed with introgressions from the wild eggplant donor parental *S. incanum* L. is available (Gramazio et al., 2017). *Solanum incanum* can tolerate long periods without irrigation (Plazas et al., 2022), and it is known that some of these ILs carry candidate genes for drought tolerance related to osmolyte and hormone biosynthesis (Gramazio et al., 2017). This set of ILs has already allowed the detection of QTLs for plant, flower, and fruit characteristics (Mangino et al., 2020), fruit morphology (Mangino et al., 2021), fruit composition (Rosa-Martinez et al., 2022a) and nitrogen fertilization (Rosa-Martinez et al., 2022b), demonstrating that this new genetic resource is of great interest for eggplant breeding. Therefore, we hypothesize that the evaluation of this set of ILs for tolerance to water stress may allow identifying genomic regions introgressed from *S. incanum* into the genetic background of eggplant, conferring improved drought tolerance.

Here, we evaluated ILs of eggplant developed with introgressed genetic material from the wild species *S. incanum* and compared them with their respective parents under conditions of proper and limited water availability. The study assessed the effects of water stress on various growth and biomass production traits and several key biochemical parameters. Our findings provide valuable insights for the incorporation of *S. incanum* and its ILs into the breeding pipeline for drought tolerance in eggplant.

## 2. Materials and methods

### 2.1 Plant material

The plant material used consisted of nine eggplant introgression lines (ILs) in the genetic background of accession AN-S-26, each line representing a single introgression from the wild eggplant relative *S. incanum* (accession MM577). The nine ILs (SMI-1-2, SMI-2-9, SMI-3-2, SMI-4-3, SMI-5-2, SMI-6-1, SMI-7-2, SMI-8-4 and SMI-12-1) carry *S. incanum* introgressions in chromosomes 1, 2, 3, 4, 5, 6, 7, 8 and 12, respectively, with 71.6% coverage of the *S. incanum* genome (Figure 1). Further details on the parents and ILs and the development of the ILs can be found in Gramazio et al. (2017), whereas phenotypic characteristics can be found in Mangino et al. (2020, 2021) and Rosa-Martínez et al. (2022a,b).

**Figure 1.**
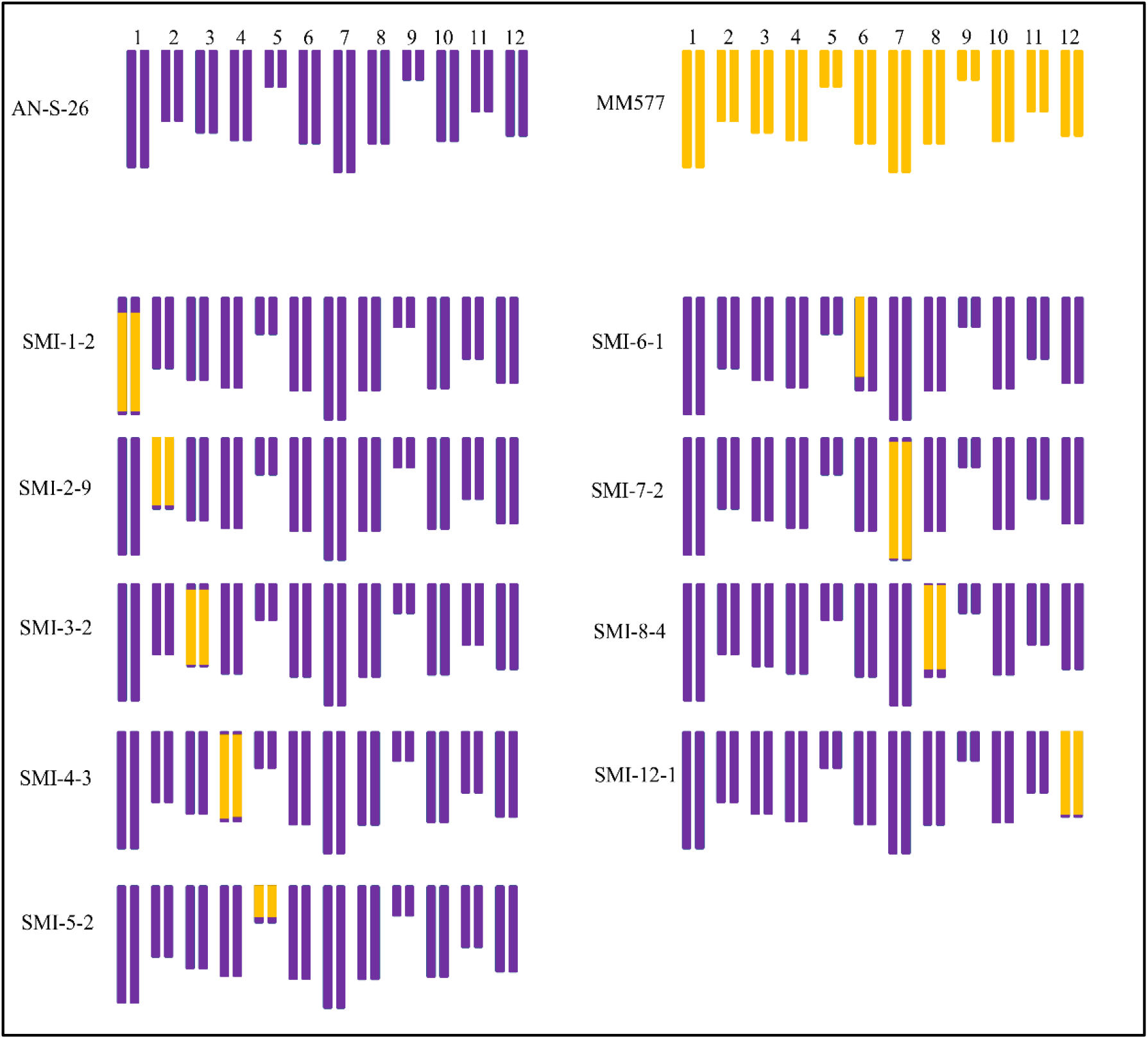
Genotypes of the ILs used. The genetic background of *S. melongena* (AN-S-26) is represented in purple, and the introgressions of *S. incanum* (MM577) are represented in yellow. The number of each chromosome is shown above.

### 2.2 Growing conditions

The seeds of the nine introgression lines (ILs) and their parental accessions AN-S-26 and MM577, were germinated in Petri dishes according to the protocol described by Ranil et al. (2015). The seedlings were then transplanted into trays filled with Humin substrate N3 (Klasmann-Deilmann, Germany) and grown in a greenhouse with controlled temperature (maximum 30ºC and minimum 15ºC). After 34 days, the plants were transferred to individual 1.3 L pots filled with the same substrate and watered every two days to field capacity (FC).

Water stress treatment began 55 days after germination when plants had five developed true leaves. This consisted of water replacement up to FC for control plants and water replacement only up to 30% of FC for the water stress treatment (Figure 2). The gravimetric water content at field capacity (*θ*_FC_) for each pot was calculated as described in Rolando et al. (2015). The pot was weighted (PW) and then filled with the equivalent of 144 g of dry substrate and reweighed (DW). Subsequently, the pots were watered to saturation and covered at the top with aluminium foil to prevent evaporation. After 48 hours, when no drainage was observed, they were weighed again (SCW), and the amount of water retained by the substrate was calculated. In this way, the *θ*_FC_ was estimated as:

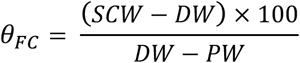

**Figure 2.**
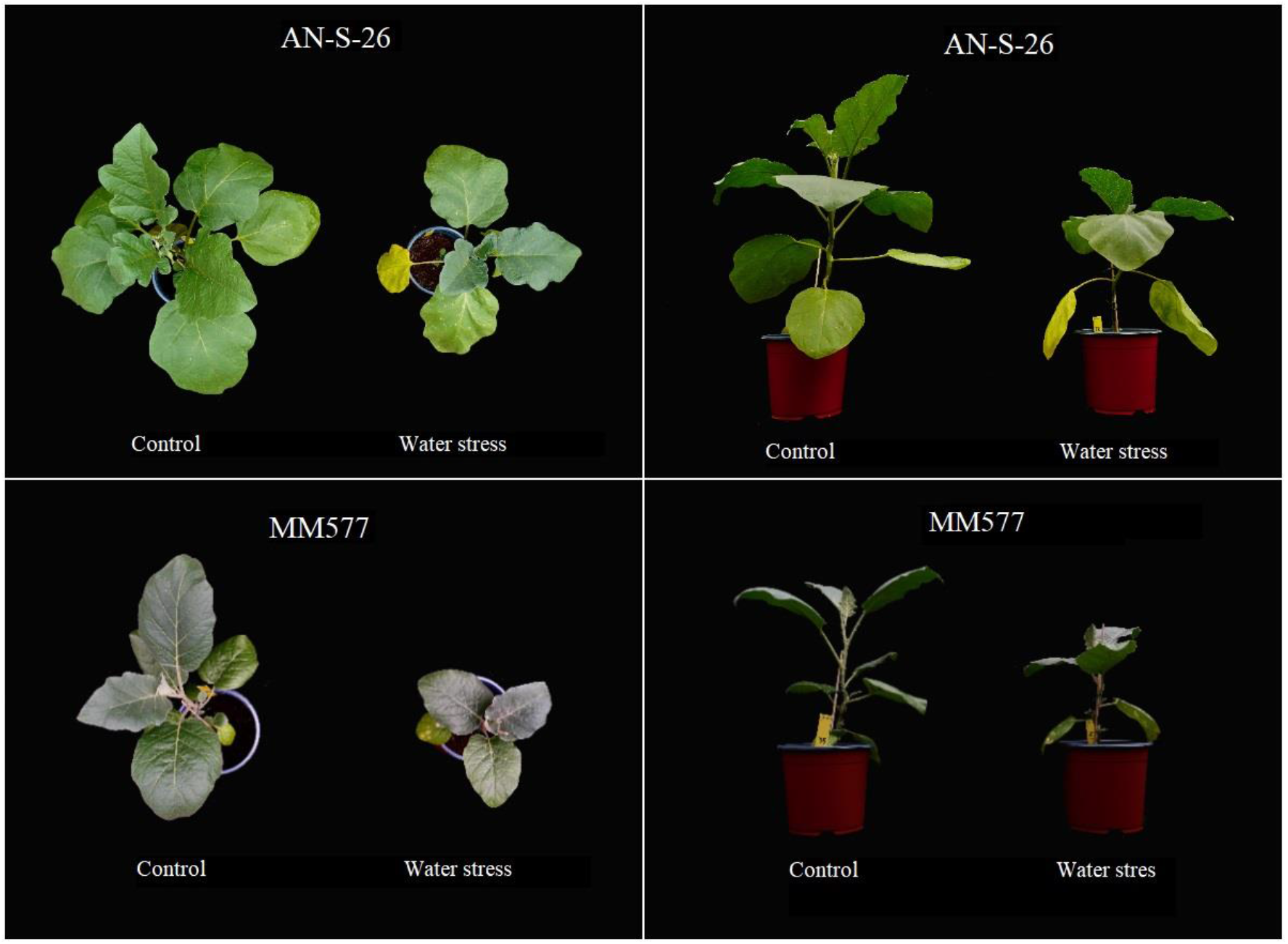
Parental of the ILs in irrigated at 100% (Control) and 30% (Water stress) field capacity at the end of the experiment. *S. melongena* AN-S-26 is placed at the top, and *S. incanum* MM577 at the bottom. Upper (left panels) and lateral (right panels) views of the plants are shown.

The amount of water in the pot at this point was considered 100% FC, and from this, the pot weight was calculated for a moisture content of 30% FC for the water stress treatment.

Five plants per genotype were selected for each treatment, except for the parental AN-S-26, where fifteen plants per treatment were used. To minimise the effects of microclimatic differences within the greenhouse, potted plants were randomly distributed and moved every two days. From the beginning of the water treatments, the plants were watered daily by measuring the weight of each pot and adding the corresponding water until the pot weight reached 100% or 30% of the substrate field capacity (FC). The treatments were continued for 14 days, after which the plants were evaluated for morphological and biochemical characteristics.

### 2.3 Morphological evaluation and water content

At the end of the experiment, plant fresh weight (FW), leaf fresh weight (FW leaf), stem length (SL) and the number of leaves (Lno) were measured. The fresh material of each plant organ was then dried at 65ºC for 72 hours to obtain aerial dry weight (DW aerial), leaf dry weight (DW leaf), stem dry weight (DW stem) and root dry weight (DW root). These data were used to calculate the leaf water content (LWC), expressed as the percentage of water contained in the samples [1]. The water use efficiency (WUE) was calculated from the water applied during the experimental period (Irrigation) and the FW [2].

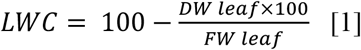

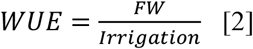

### 2.4 Biochemical analyses

For all biochemical analyses, fresh leaf samples were taken from each plant. The concentrations of photosynthetic pigments, chlorophyll (Chl) and carotenoids (Caro), were measured according to the method described by Lichtenthaler and Walllburn (1983). A 50 mg sample of fresh leaves was extracted with 1 ml of 80% (v/v) acetone and left for 24 hours in the dark with stirring. After centrifugation, the supernatant was used to measure absorbance at 663, 646 and 470 nm. The Chl and Caro concentrations were then calculated using the equations of Lichtenthaler and Wallburn (1983).

Proline (Pro) was quantified by the ninhydrin-acetic acid method (Bates et al., 1973). A 0.10 g sample of fresh leaves was ground and mixed with 1 ml of sulphosalicylic acid solution (3%, w/v). The mixture was treated with acid ninhydrin and incubated in a water bath at 98ºC for 1 hour, then cooled on ice for 10 minutes. Finally, proline was extracted with toluene, and the absorbance of the organic phase was measured at 520 nm, using toluene as the blank.

For the determination of malondialdehyde (MDA), total flavonoids (TF) and total phenolic compounds (TPC), an extract was prepared with 80% (v/v) methanol and 0.1 g of fresh leaves. For the quantification of MDA, 200 μl of the extract was mixed with 0.5% thiobarbituric acid and 20% trichloroacetic acid; another 200 μl of the extract, mixed with 20% trichloroacetic acid only, was used as the corresponding control. The samples were heated at 95ºC for 15 minutes and then cooled on ice to stop the reaction. Absorbance at 440, 532 and 600 nm was measured to calculate concentrations using the equations described in Hodges et al. (1999).

Total phenolic compounds (TPC) content was measured by reacting the methanol extract with the Folin-Ciocalteu reagent (Blainski et al., 2013). Na_2_CO_3_ was added, and the mixture was incubated in the dark for 90 min at room temperature. The absorbance was measured at 765 nm, and the equivalent concentration was calculated using gallic acid as a standard.

Total flavonoids (TF) were quantified following the method used by Zhishen et al. (1999). NaNO_2_ was added to the methanol extract, followed by AlCl_3_ and NaOH, and the absorbance was measured at 510 nm. Catechin was used as a standard to calculate the equivalent concentration.

### 2.5 Data analysis

For each trait, data collected within each combination of genotype and treatment were subjected to a two-factor analysis of variance (ANOVA) to determine differences between genotypes, treatments, and the interaction between treatments and genotypes. Treatment means within each genotype were compared using the Student-Newman-Keuls (SNK) multiple comparison test at a significance level of *p*<0.05.

Pearson’s correlation coefficients were used to perform correlation tests within each treatment to examine the degree of association between variables. Principal component analysis (PCA) was also performed to analyse the relationship between the measured traits and genotypes.

Analyses were performed using the statistical software R 4.2.1 (Team R. C., 2013) and the packages *psych* (Revelle, 2017) and *corrplot* (Wei and Simko, 2017) for correlations and *stats* and *ggplot2* (Wickham, 2009) for PCA.

### 2.6 QTL detection

The ILs data were analysed to determine the presence of QTLs in the genomic region of introgression. A Dunnett’s test at a significance level of *p*<0.05 was performed to compare each IL with the cultivated parent (AN-S-26) for each trait. A QTL was considered stable and of high effect if it complied with the following conditions: i) significant (*p*<0.05) in both conditions; ii) same sign (positive or negative) in both conditions; and iii) caused a change (increase or decrease) of more than 25% compared to AN-S-26 in at least one of the two conditions. In addition, QTLs were determined for the percentage of change of each trait when grown under well-watered conditions and under water stress, and those that caused an increase or decrease of more than 25% compared to AN-S-26 were selected.

## 3. Results

### 3.1 Analysis of variance

The ANOVA performed on the nine ILs and their parents showed significant differences (*p*<0.05) for all traits in at least one of the factors examined (treatment, genotype, and treatment × genotype) (Table 1). Highly significant differences (*p*<0.01) were found within treatment for all traits except for TPC and TF, which showed no significant differences. For the factor genotype, 13 of the 14 traits showed highly significant differences (*p*<0.01), while for the treatment × genotype interaction, only seven of the 14 traits showed significant differences (Table 1). According to the contribution to the total sum of squares (SS), the treatment was the source of variation that contributed most to the total variance of the growth and proline content parameters (Lno, SL and DW), whereas the genotype had a greater contribution to LWC and WUE. The interaction effect was not the main contributor to any of the traits, whereas the residual effect had a greater contribution to the biochemical parameters Chl, Caro, MDA, TPC and TF (Table 1).

**Table 1.** Analysis of variance for treatment, genotype, and interaction treatment by genotype, as well as the residual for the parameters evaluated. The contribution to the total sum of squares (SS; %) and the *p*-value are shown. Abbreviations: leaf number (Lno), stem length (SL), aerial dry weight (DW aerial), leaf dry weight (DW leaf), stem dry weight (DW stem), root dry weight (DW root), leaf water content (LWC), water use efficiency (WUE), chlorophyll (Chl), carotenoids (Caro), proline (Pro), malondialdehyde (MDA), total phenolic compounds (TPC), total flavonoids (TF).

| Trait | Treatment |  | Genotype |  | Treatment x Genotype |  | Residual |
| --- | --- | --- | --- | --- | --- | --- | --- |
| | SS (%) | $p$-value | SS (%) | $p$-value | SS (%) | $p$-value | SS (%) |
| Lno | 67.1 | <0.0001 | 11.7 | <0.0001 | 2.6 | 0.1523 | 18.6 |
| SL (cm) | 49.7 | <0.0001 | 39.9 | <0.0001 | 5.2 | <0.0001 | 5.2 |
| DW aerial (g) | 63.3 | <0.0001 | 23.7 | <0.0001 | 4.5 | <0.0001 | 8.5 |
| DW leaf (g) | 62.2 | <0.0001 | 22.2 | <0.0001 | 4.0 | 0.0003 | 11.6 |
| DW stem (g) | 50.2 | <0.0001 | 33.7 | <0.0001 | 6.6 | <0.0001 | 9.4 |
| DW root (g) | 34.4 | <0.0001 | 27.3 | <0.0001 | 9.7 | 0.0004 | 28.6 |
| LWC (%) | 2.9 | 0.0015 | 61.8 | <0.0001 | 6.3 | 0.0175 | 29.2 |
| WUE (g FW ml<sup>-1</sup>) | 12.0 | <0.0001 | 66.5 | <0.0001 | 4.4 | 0.0490 | 17.1 |
| Chl (mg g<sup>-1</sup> DW) | 7.8 | 0.0002 | 26.2 | <0.0001 | 9.7 | 0.0658 | 56.3 |
| Caro (mg g<sup>-1</sup> DW) | 4.1 | 0.0084 | 27.1 | <0.0001 | 9.2 | 0.1042 | 59.6 |
| Pro (μmoles g<sup>-1</sup> DW) | 39.0 | <0.0001 | 13.4 | 0.0002 | 10.3 | 0.0027 | 37.2 |
| MDA (μmoles g<sup>-1</sup> DW) | 19.7 | <0.0001 | 14.6 | 0.0081 | 5.6 | 0.4610 | 60.2 |
| TPC (mg eq. GA g<sup>-1</sup> DW) | 0.5 | 0.3352 | 32.5 | <0.0001 | 8.7 | 0.1223 | 58.3 |
| TF (mg eq. C g<sup>-1</sup> DW) | 0.2 | 0.5625 | 30.2 | <0.0001 | 3.1 | 0.8873 | 66.4 |

### 3.2 Growth and water content

In both parents and all ILs, the growth parameters Lno and SL decreased significantly when grown under water stress conditions (Table 2). The mean values obtained in the cultivated parent (AN-S-26) were higher than in the wild parent (MM577), but the percentage of decrease was lower in the latter. The ILs presented Lno and SL average values intermediate between those of the two parents (Table 2).

**Table 2.** Effect of water stress (WS) in ILs and their parents (AN-S-26 and MM577) for control and water stress conditions and the percentage of change in the WS treatment with respect to the control. Abbreviations: leaf number (Lno), stem length (SL), aerial dry weight (DW aerial), leaf dry weight (DW leaf), stem dry weight (DW stem), root dry weight (DW root), leaf water content (LWC) and water use efficiency (WUE).

| Genotype | Treatment | Lno | SL<br>(cm) | DW<br>aerial<br>(g) | DW leaf<br>(g) | DW stem<br>(g) | DW root<br>(g) | LWC<br>(%) | WUE<br>(g FW ml<sup>-1</sup>) |
| --- | --- | --- | --- | --- | --- | --- | --- | --- | --- |
| AN-S-26 | Control | 12.7 | 34.4 | 7.25 | 4.82 | 2.35 | 2.93 | 90.6 | 0.0175 |
|  | WS | 7.3 | 23.6 | 3.79 | 2.52 | 1.27 | 1.43 | 89.9 | 0.0203 |
|  | Change (%) | -42.4%*** | -31.6%*** | -47.8%*** | -45.8%*** | -51.1%*** | -51.1%*** | -0.7%* | 15.7%*** |
| MM577 | Control | 9.4 | 20.9 | 2.77 | 1.95 | 0.82 | 0.99 | 86.5 | 0.0077 |
|  | WS | 5.8 | 14.5 | 1.58 | 1.01 | 0.57 | 0.86 | 84.5 | 0.0092 |
|  | Change (%) | -38.3%*** | -30.6%*** | -42.9%* | -48.4%** | -29.9% | -13.3% | -2.3%* | 19.5% |
| SMI-1-2 | Control | 10.0 | 24.5 | 4.82 | 3.38 | 1.35 | 2.02 | 91.8 | 0.0150 |
|  | WS | 6.5 | 15.9 | 2.75 | 2.07 | 0.67 | 1.06 | 90.3 | 0.0193 |
|  | Change (%) | -35.0%*** | -35.2%*** | -43.1%*** | -38.7%*** | -50.3%** | -47.8%* | -1.6%*** | 29.4%** |
| SMI-2-9 | Control | 10.2 | 26.0 | 6.00 | 4.20 | 1.74 | 2.10 | 90.4 | 0.0153 |
|  | WS | 5.8 | 15.7 | 2.53 | 1.82 | 0.71 | 1.01 | 90.0 | 0.0169 |
|  | Change (%) | -43.1%*** | -39.6%*** | -57.8%*** | -56.7%*** | -59.2%*** | -52.1%* | -0.5% | 10.2% |
| SMI-3-2 | Control | 10.2 | 26.0 | 6.09 | 4.06 | 1.94 | 2.02 | 90.5 | 0.0149 |
|  | WS | 5.8 | 15.7 | 2.91 | 2.06 | 0.84 | 1.23 | 88.4 | 0.0146 |
|  | Change (%) | -44.9%*** | -47.5%*** | -52.2%*** | -49.2%*** | -56.5%*** | -39.1%* | -2.4%** | -2.2% |
| SMI-4-3 | Control | 10.8 | 31.9 | 7.93 | 5.38 | 2.39 | 4.34 | 88.8 | 0.0158 |
|  | Water stress | 5.6 | 18.7 | 3.80 | 2.70 | 1.09 | 1.72 | 88.1 | 0.0169 |
|  | Change (%) | -48.1%*** | -41.4%*** | -52.1%*** | -49.8%*** | -54.3%*** | -60.4%** | -0.8% | 6.7% |
| SMI-5-2 | Control | 8.6 | 18.5 | 7.45 | 5.75 | 1.70 | 3.02 | 87.8 | 0.0157 |
|  | WS | 5.8 | 14.9 | 3.72 | 2.79 | 0.93 | 1.77 | 88.0 | 0.0194 |
|  | Change (%) | -32.6%** | -19.5%** | -50.1%** | -51.5%** | -45.3%*** | -41.6%* | 0.3% | 23.2%* |
| SMI-6-1 | Control | 10.4 | 20.6 | 6.36 | 4.94 | 1.42 | 2.75 | 88.7 | 0.0150 |
|  | WS | 6.0 | 14.8 | 3.13 | 2.40 | 0.73 | 2.02 | 88.0 | 0.0187 |
|  | Change (%) | -42.3%*** | -28.2%*** | -50.8%*** | -51.5%*** | -48.5%*** | -26.3% | -0.8% | 24.8%** |
| SMI-7-2 | Control | 11.6 | 34.2 | 8.01 | 4.83 | 2.96 | 4.58 | 89.3 | 0.0172 |
|  | WS | 6.8 | 19.4 | 3.68 | 2.47 | 1.19 | 1.70 | 89.0 | 0.0188 |
|  | Change (%) | -41.4%*** | -43.3%*** | -54.1%*** | -48.8%*** | -59.7%*** | -63.0%*** | -0.3% | 9.6% |
| SMI-8-4 | Control | 11.4 | 36.3 | 8.85 | 5.64 | 3.04 | 3.57 | 89.2 | 0.0169 |
|  | WS | 6.4 | 23.0 | 3.87 | 2.42 | 1.45 | 1.95 | 90.6 | 0.0210 |
|  | Change (%) | -43.9%*** | -36.6%*** | -56.2%*** | -57.1%*** | -52.2%*** | -45.4%** | 1.5% | 24.5%* |
| SMI-12-1 | Control | 10.8 | 30.8 | 7.29 | 4.77 | 2.31 | 2.53 | 89.9 | 0.0155 |
|  | WS | 4.4 | 20.6 | 3.03 | 1.93 | 1.06 | 1.55 | 90.1 | 0.0158 |
|  | Change (%) | -59.3%*** | -33.1%*** | -58.4%*** | -59.5%*** | -54.1%*** | -38.8%* | 0.2% | 2.2% |

\*, \*\*, \*\*\* indicate significance in $p < 0.05$ , $< 0.01$ or $< 0.001$ , respectively (SNK Multiple comparison test).

Water stress significantly affected DW, with most genotypes showing losses of more than 50% of biomass compared to the control (Table 2). Aerial DW, leaf DW and stem DW decreased in all ILs and the cultivated parent; however, in MM577 no significant differences were observed in stem DW due to water stress. Meanwhile, root DW decreased at lower irrigation levels for AN-S-26 and eight of the nine ILs, whereas no significant treatment effects were observed for MM577 and SMI-6-1. A lower water stress effect was observed in the MM577 parent, the average DW values in leaves, stems and roots in both conditions were much lower than those of AN-S-26 and the ILs.

LWC did not display a wide range of values, with most of the genotypes evaluated showing no significant differences between control and water stress conditions (Table 2). However, both parents, AN-S-26 and MM577, showed a decrease in LWC in response to water stress, but this was not observed in most of the ILs, where only SMI-1-2 and SMI-2-9 decreased under water stress conditions. On the other hand, water stress increased WUE in the cultivated parent AN-S-26 and four of the nine ILs, reaching an increase of up to 29.4% in SMI-1-2. The wild parent did not increase WUE under stress conditions and showed much lower average values than the cultivated parent and the ILs (Table 2).

### 3.3 Biochemical analyses

Water treatments affected photosynthetic pigments, with leaf Chl and Caro decreasing in response to water stress in some genotypes (Table 3). In this case, the parental MM577 was affected by stress, decreasing its Chl and Caro content by 38.4% and 35.3%, respectively; on the contrary, pigment concentrations did not show significant differences between the two treatments in AN-S-26. On the other hand, SMI-1-2 and SMI-12-1 showed a significant decrease in Chl, and SMI-1-2 also showed a decrease in Caro. Thus, no significant negative effects of water stress on photosynthetic pigments were observed for seven of the nine ILs (Table 3).

**Table 3.** Effect of water stress on biochemical parameters in ILs and their parents (AN-S-26 and MM577). Abbreviations: chlorophyll (Chl), carotenoids (Caro), proline (Pro), malondialdehyde (MDA), total phenolic compounds (TPC), total flavonoids (TF).

| Genotype | Treatment | Chl<br>(mg g<sup>-1</sup> DW) | Caro<br>(mg g<sup>-1</sup> DW) | Pro<br>(μmol g<sup>-1</sup> DW) | MDA<br>(nmol g<sup>-1</sup> DW) | TPC<br>(mg eq. GA g<sup>-1</sup> DW) | TF<br>(mg eq. C g<sup>-1</sup> DW) |
| --- | --- | --- | --- | --- | --- | --- | --- |
| AN-S-26 | Control | 12.5 | 5.2 | 5.8 | 672.6 | 17.5 | 15.3 |
|  | Water stress | 11.7 | 4.9 | 52.8 | 1048.3 | 19.8 | 18.8 |
|  | Change (%) | -6.4% | -6.0% | 810.3%*** | 55.9%*** | 13.2% | 23.0% |
| MM577 | Control | 13.6 | 4.8 | 55.7 | 620.9 | 6.6 | 4.7 |
|  | Water stress | 8.4 | 3.1 | 75.8 | 694.8 | 12.7 | 7.6 |
|  | Change (%) | -38.4%* | -35.3%** | 36.1% | 11.9% | 93.1%*** | 61.7%* |
| SMI-1-2 | Control | 25.7 | 9.0 | 14.8 | 1009.3 | 23.8 | 14.6 |
|  | Water stress | 14.5 | 6.1 | 48.9 | 1116.5 | 15.2 | 9.3 |
|  | Change (%) | -43.5%* | -32.4%* | 230.4%* | 10.6% | -36.4% | -36.4% |
| SMI-2-9 | Control | 20.3 | 7.2 | 19.2 | 710.6 | 12.7 | 9.0 |
|  | Water stress | 17.2 | 6.9 | 70.9 | 1024.3 | 17.6 | 11.9 |
|  | Change (%) | -15.2% | -4.0% | 269.3%* | 44.1% | 38.4%* | 32.7% |
| SMI-3-2 | Control | 18.7 | 7.0 | 7.7 | 704.3 | 19.4 | 15.4 |
|  | Water stress | 10.7 | 4.1 | 20.3 | 812.6 | 19.0 | 16.5 |
|  | Change (%) | -42.7% | -40.9% | 163.6%* | 15.4% | -2.1% | 7.3% |
| SMI-4-3 | Control | 13.5 | 5.5 | 6.0 | 595.4 | 22.1 | 18.2 |
|  | Water stress | 13.4 | 5.2 | 73.4 | 905.7 | 21.4 | 17.6 |
|  | Change (%) | -0.7% | -5.8% | 1123.3%*** | 52.1%* | -3.4% | -3.3% |
| SMI-5-2 | Control | 12.3 | 4.3 | 7.2 | 709.9 | 18.2 | 12.8 |
|  | Water stress | 11.0 | 4.5 | 92.3 | 891.4 | 14.7 | 9.7 |
|  | Change (%) | -10.1% | 5.6% | 1181.9%*** | 25.6% | -19.4% | -24.3% |
| SMI-6-1 | Control | 11.4 | 4.7 | 11.8 | 553.0 | 9.5 | 6.7 |
|  | Water stress | 9.3 | 4.0 | 35.4 | 788.5 | 11.6 | 5.2 |
|  | Change (%) | -18.0% | -15.5% | 200.0%* | 42.6% | 21.8% | -22.4% |
| SMI-7-2 | Control | 12.8 | 4.8 | 9.9 | 890.8 | 20.4 | 13.2 |
|  | Water stress | 12.1 | 4.9 | 33.7 | 1067.1 | 18.0 | 12.7 |
|  | Change (%) | -5.2% | 2.5% | 240.4%* | 19.8% | -11.8% | -3.5% |
| SMI-8-4 | Control | 13.2 | 5.0 | 3.9 | 705.6 | 18.5 | 17.3 |
|  | Water stress | 13.4 | 6.1 | 88 | 1292.5 | 22.9 | 17.6 |
|  | Change (%) | 1.6% | 21.4% | 2156.4%* | 83.2%* | 23.3% | 1.8% |
| SMI-12-1 | Control | 20.0 | 7.1 | 11.3 | 771.1 | 18.2 | 13.3 |
|  | Water stress | 12.7 | 5.4 | 100.5 | 1319.3 | 19.3 | 14.1 |
|  | Change (%) | -36.5%* | -25.1% | 789.4%** | 71.1% | 6.1% | 5.8% |

\*, \*\*, \*\*\* indicate significance in $p < 0.05$ , $< 0.01$ or $< 0.001$ , respectively (SNK Multiple comparison test).

Water stress conditions affected the Pro content of most of the evaluated genotypes (Table 3), although the difference between genotypes was not significant (Table 1). Although the Pro content of AN-S-26 increased by 810% due to water stress, the wild genotype (MM577) showed no significant differences between the two treatments. Meanwhile, all ILs significantly increased Pro content under water stress conditions, with very different percentages, ranging from 164% in SMI-6-1 to 2156% in SMI-8-4.

Although the ANOVA showed a significant effect of the water stress treatment on MDA contents (Table 1), most genotypes did not show significant differences in their values. Thus, the only genotypes showing significant differences in MDA concentrations were the cultivated parent (AN-S-26), with an increase of 56%, and the ILs SMI-4-3 and SMI-8-4, with increases of 52% and 83%, respectively (Table 3).

For non-enzymatic antioxidants, significant differences between genotypes were found only in the parental lines MM577 and SMI-2-9 for TPC, which increased by 93 and 38%, respectively. MM577 showed a significant increase of 62% in TF, but the ILs did not change their levels when grown under water stress conditions (Table 3).

### 3.4 Correlation and Principal Component Analyses

Correlation coefficients were calculated separately for control and water stress conditions for the 14 traits evaluated (Figure 3). Under control conditions, high positive correlations (greater than 0.80) were observed between growth traits such as aerial DW with leaf DW and stem DW, and stem DW with LS. WUE also showed positive correlations, greater than 0.5, for SL and all DW traits. For biochemical traits, high positive correlations were found between Chl and Caro, and TPC and TF. Regarding the correlations between growth and biochemical traits, a negative relationship was observed between proline and aerial DW, leaf DW and WUE; on the other hand, TPC was positively correlated with stem DW, and photosynthetic pigments (Chl and Caro) with LWC (Figure 3).

**Figure 3.**
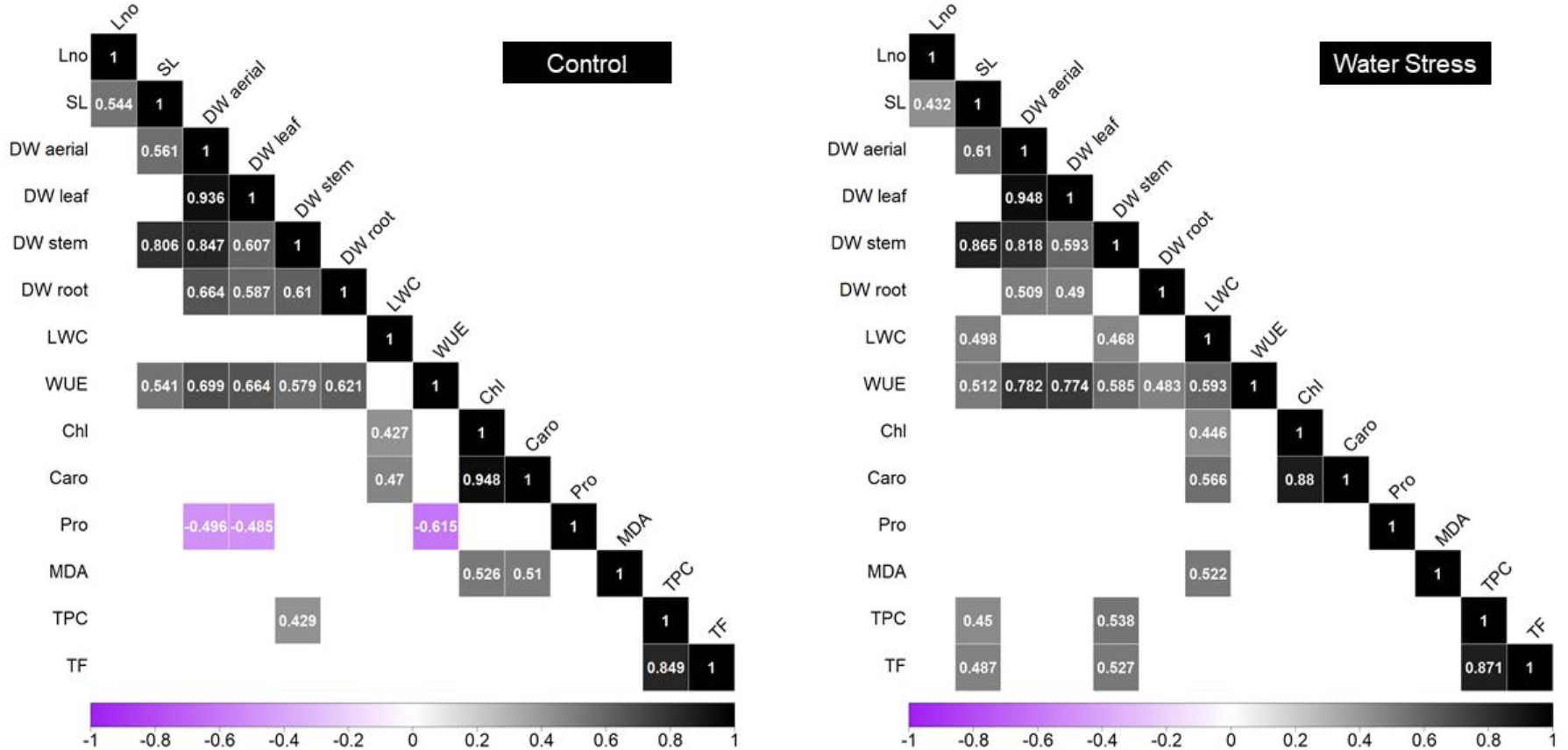
Correlation matrix coefficients for control and water stress in ILs and their parents (AN-S-26 and MM577). Only statistically significant correlations (*p*-value < 0.05) are shown. Abbreviations: leaf number (Lno), stem length (SL), aerial dry weight (DW aerial), leaf dry weight (DW leaf), stem dry weight (DW stem), root dry weight (DW root), leaf water content (LWC), water use efficiency (WUE), chlorophyll (Chl), carotenoids (Caro), proline (Pro), malondialdehyde (MDA), total phenolic compounds (TPC) and total flavonoids (TF).

Under water stress conditions, none of the evaluated traits showed significant negative correlations (Figure 3). Significant positive correlations greater than 0.80 were observed between the same traits as those observed under control conditions, where positive correlations were found between the growth traits aerial DW, leaf DW and stem DW, and between stem DW and LS; also, regarding biochemical traits, between Chl and Caro, and between TPC and TF (Figure 3). In this treatment, WUE showed positive correlations greater than 0.5 with SL and aerial DW, leaf DW, stem DW and LWC. Amongst the significant correlations between growth and biochemical traits under water stress conditions, positive correlations were observed between non-enzymatic antioxidants (TPC and TF) and stem growth (SL and stem DW), and between photosynthetic pigments (Chl and Caro) and LWC (Figure 3).

A PCA was performed to determine the relationships between the traits measured under control and water stress conditions and to associate them with the genotypes studied. The first (PC1) and second (PC2) components accounted for 47.8% and 24.5% of the total variation, respectively (Figure 4). PC1 was positively correlated with growth traits and negatively correlated with biochemical markers of water stress (MDA and Pro). PC2 was positively correlated with biochemical parameters, LWC and WUE, and negatively correlated with DW and Lno (Figure 4). All ILs and the recurrent parental *S. melongena* AN-S-26 evaluated under control conditions had positive values for PC1 (Figure 4). However, the donor wild parental *S. incanum* MM577, and all genotypes evaluated under water stress conditions, had negative values for PC1. For PC2, most of the genotypes under control conditions were grouped in the lower part with negative values, whereas most of the genotypes under water stress were grouped in the upper part with positive values (Figure 4). The three genotypes with the lowest values for PC2 are the same in both conditions (MM577, SMI-6-1 and SMI-5-2). The ellipses grouping the genotypes with 95% significance within each treatment had a minimal overlap, although a wider distribution was observed in the control treatment, indicating a greater variability of the traits in this condition (Figure 4).

**Figure 4.**
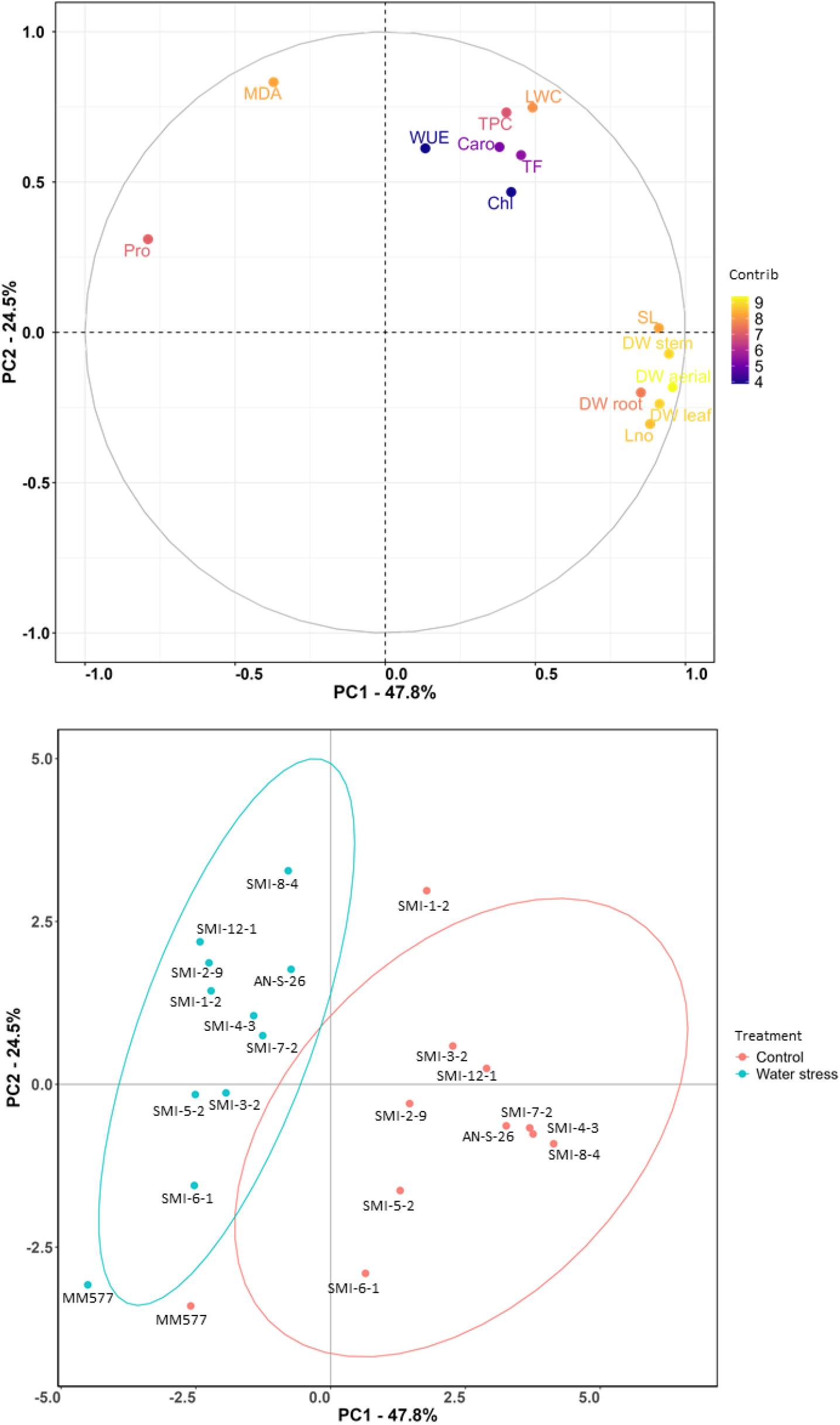
Loading plot (above) and score plot (below) of the principal component analysis (PCA) for the 9 ILs and their parents (AN-S-26 and MM577), based on the first two principal components. The first and second components (PC1 and PC2) represent 48.6% and 23.4% of the variation, respectively. The ellipses in the score plot group the genotypes for each treatment with a confidence level of 95%. Abbreviations: leaf number (Lno), stem length (SL), aerial dry weight (DW aerial), leaf dry weight (DW leaf), stem dry weight (DW stem), root dry weight (DW root), leaf water content (LWC), water use efficiency (WUE), chlorophyll (Chl), carotenoids (Caro), proline (Pro), malondialdehyde (MDA), total phenolic compounds (TPC) and total flavonoids (TF).

### 3.5 QTL detection

A total of 19 stable and high-effect QTLs were detected in both environments, with a percentage of variation greater than 25% in at least one of the growth conditions (Table 4). For Lno, three QTLs were located on chromosomes 3, 5 and 12; for SL, four QTLs were detected on chromosomes 1, 2, 5 and 6. For DW, nine QTLs were found present in both treatments, most of them affecting the stem DW (Table 4). Three of them were co-located on chromosome 1 (*dwa1, dwl1* and *dws1*), two on chromosome 2 (*dwa2* and *dws2*) and four others, affecting only the stem, on chromosomes 3, 5, 6 and 8. One QTL was found for WUE, located on chromosome 3. For biochemical traits, two QTLs were found, one for chlorophyll content (chl2), located on chromosome 2, and one for TPC, located on chromosome 6 (Table 4).

**Table 4.** List of stable and high-effect QTLs detected in the eggplant IL population, present in both conditions and generating an increase or decrease of more than 25% when in homozygous condition in at least one of the two treatments, with respect to the parental AN-S-26. The allelic effect refers to the additive effect caused by the *S. incanum* allele. Abbreviations: leaf number (Lno), stem length (SL), aerial dry weight (DW aerial), leaf dry weight (DW leaf), stem dry weight (DW stem), water use efficiency (WUE) chlorophyll (Chl) and total phenolic compounds (TPC).

| Trait | QTL | Chr. | Position (Mb) | Control |  |  | Water Stress |  |  |
| --- | --- | --- | --- | --- | --- | --- | --- | --- | --- |
|  |  |  |  | Increase Over AN-S-26 (%) | Allelic Effect (Units) | Significance | Increase Over AN-S-26 (%) | Allelic Effect (Units) | Significance |
| Lno | *lno3* | 3 | 7.5-86.6 | -22.8 | -1.5 | ** | -26.0 | -1.0 | ** |
| Lno | *lno5* | 5 | 0.0-36.4 | -32.3 | -2.1 | *** | -20.5 | -0.8 | ** |
| Lno | *lno12* | 12 | 2.4-97.0 | -15.0 | -0.9 | * | -39.7 | -1.5 | *** |
| SL | *sl1* | 1 | 20.8- 134.0 | -28.8 | -5.0 | *** | -32.6 | -3.9 | *** |
| SL | *sl2* | 2 | 0.0-77.5 | -24.4 | -4.2 | *** | -33.5 | -4.0 | *** |
| SL | *sl5* | 5 | 0.0-36.4 | -46.2 | -8.0 | *** | -36.9 | -4.4 | *** |
| SL | *sl6* | 6 | 4.7-92.2 | -40.1 | -13.8 | *** | -37.3 | -8.8 | *** |
| DW aerial | *dwa1* | 1 | 20.8- 134.0 | -33.4 | -1.2 | *** | -27.4 | -0.5 | *** |
| DW aerial | *dwa2* | 2 | 0.0-77.5 | -17.3 | -0.6 | * | -33.2 | -0.6 | *** |
| DW leaf | *dwl1* | 1 | 20.8- 134.0 | -29.9 | -0.7 | ** | -17.5 | -0.2 | * |
| DW stem | *dws1* | 1 | 20.8- 134.0 | -42.5 | -0.5 | *** | -47.2 | -0.3 | *** |
| DW stem | *dws2* | 2 | 0.0-77.5 | -25.9 | -0.3 | ** | -44.1 | -0.3 | *** |
| DW stem | *dws3* | 3 | 7.5-86.6 | -17.3 | -0.2 | * | -33.9 | -0.2 | *** |
| DW stem | *dws5* | 5 | 0.0-36.4 | -27.6 | -0.3 | ** | -26.8 | -0.2 | *** |
| DW stem | *dws6* | 6 | 4.7-92.2 | -39.5 | -0.9 | *** | -42.5 | -0.5 | *** |
| DW stem | *dws8* | 8 | 1.8-99.6 | 29.4 | 0.3 | ** | 14.2 | 0.1 | ** |
| WUE | *wue3* | 3 | 7.5-86.6 | -14.9 | -0.0013 | ** | -28.1 | -0.0028 | *** |
| Chl | *chl2* | 2 | 0.0-77.5 | 61.5 | 3.9 | * | 46.3 | 2.7 | *** |
| TPC | *tpc6* | 6 | 4.7-92.2 | -45.8 | -8.0 | * | -41.5 | -8.2 | ** |

\*, \*\*, \*\*\* indicate significance in $p < 0.05$ , $< 0.01$ or $< 0.001$ , respectively (Dunnett test).

All the QTLs shown in Table 4 have negative effects on plant development except for two of them, *dws8*, which causes an increase of 29.4% and 14.2% over the parental *S. melongena* AN-S-26 under control and water stress conditions, respectively, and *chl2*, which causes an increase of 61.5% and 46.3% over AN-S-26 under control and water stress conditions, respectively.

For the traits evaluated, a total of 22 QTLs were found with an effect greater than 25% with respect to the parental AN-S-26 for the percentage of change in the traits between the control and water stress treatments (Table 5). For growth parameters, one QTL was detected for the percentage change in Lno on chromosome 12, and four for SL located on chromosomes 3, 4, 5 and 7. For DW, three QTLs were detected on chromosomes 2 and 7, related to the stem, and on chromosome 6, related to the root. For LWC variation, two QTLs were found on chromosomes 3 and 12; for WUE, three QTLs were found on chromosomes 1, 3 and 12 (Table 5). Five QTLs were also found to be related to photosynthetic pigments, one on chromosome 1 for chlorophyll, two co-localised on chromosome 3 (*chl3%* and *caro3%*), and two co-localised on chromosome 8 (*chl8%* and *caro8%*), affecting chlorophyll and carotenoids. A QTL was found on chromosome 8 for the percentage change in Pro content and on chromosome 1 for TPC (Table 5).

**Table 5.** List of QTLs detected in the eggplant IL population, for the percentage change between the control and water stress conditions, which represents an increase of more than 25% with respect to the AN-S-26 parental. Abbreviations: leaf number (Lno), stem length (SL), stem dry weight (DW stem), leaf water content (LWC), water use efficiency (WUE), chlorophyll (Chl), carotenoids (Caro), proline (Pro) and total phenolic compounds (TPC).

| Trait | QTL | Chr. | Position (Mb) | Increase Over AN-S-26 (%) | Allelic Effect (%) | Significance |
| --- | --- | --- | --- | --- | --- | --- |
| Lno | *lno12%* | 12 | 2-97 | -39.9 | -8.5 | *** |
| SL | *sl3%* | 3 | 8-87 | -50.3 | -8.0 | * |
| SL | *sl4%* | 4 | 4-99 | -31.0 | -4.9 | * |
| SL | *sl5%* | 5 | 0-36 | 38.6 | 6.1 | *** |
| SL | *sl7%* | 7 | 5- 140 | -37.0 | -5.9 | ** |
| DW stem | *dws2%* | 2 | 0-78 | -29.0 | -6.7 | *** |
| DW stem | *dws7%* | 7 | 5-140 | -30.3 | -7.0 | *** |
| DW root | *dwr6%* | 6 | 5-92 | 48.3 | 24.7 | * |
| LWC | *lwc3%* | 3 | 8-87 | -242.9 | -0.9 | ** |
| LWC | *lwc12%* | 12 | 2- 97 | 128.6 | 0.5 | *** |
| WUE | *wue1%* | 1 | 21- 134 | 86.1 | 6.8 | * |
| WUE | *wue3%* | 3 | 8-87 | -86.1 | -6.8 | ** |
| WUE | *wue12%* | 12 | 2- 97 | -85.4 | -6.8 | * |
| Chl | *chl1%* | 1 | 21- 134 | -579.7 | -18.6 | * |
| Chl | *chl3%* | 3 | 8-87 | -567.2 | -18.2 | ** |
| Chl | *chl8%* | 8 | 2-100 | 125.0 | 4.0 | * |
| Caro | *caro3%* | 3 | 8-87 | -581.7 | -17.5 | ** |
| Caro | *caro8%* | 8 | 2-100 | 455.0 | 13.7 | * |
| Pro | *pro8%* | 8 | 2-100 | 165.6 | 676.5 | *** |
| TPC | *tpc1%* | 1 | 21- 134 | -375.8 | -24.8 | ** |

\*, \*\*, \*\*\* indicate significance in $p < 0.05$ , $< 0.01$ or $< 0.001$ , respectively (Dunnett test).

Of the QTLs detected for percentage change, seven have a positive value, indicating a smaller decrease or increase in the value of the trait due to water stress compared to the parent AN-S-26. Thirteen other QTLs have a negative value, indicating that the trait had a higher decrease in percentage than AN-S-26 due to water stress (Table 5).

## 4. Discussion

Due to the impacts of climate change on agriculture, developing new cultivars that can withstand conditions of low water availability is critical (Snowdon et al., 2020). In this regard, wild relatives of crops often show increased resilience and grow in environments with limited resources (Prohens et al., 2017; Egea et al., 2018; Iseki et al., 2018). The introgression of abiotic stress resistance genes from wild species has made it possible to obtain cultivated genotypes with greater tolerance to water stress conditions in several species (Honsdorf et al., 2014; Pino et al., 2013; Souter et al., 2017). In this study, for the first time, eggplant lines with introgressions from the wild relative *S. incanum* were evaluated for their performance under water stress conditions. This species is naturally distributed from North Africa and the Middle East to Pakistan, where it is often found in arid climates (Aubriot and Knapp, 2022).

Our study, along with others, has found that water stress negatively impacts biomass production in eggplant (Çolak et al., 2015; Delfin et al., 2021; Plazas et al., 2019; Semida et al., 2021). Previous research has shown that using wild species to generate inter-specific lines (ILs) in tomato (Poudyal et al., 2017) and pearl millet (Sharma et al., 2020) can improve biomass production under drought conditions. However, in our case study, introgressions of *S. incanum* into *S. melongena* background exhibited mostly negative effects on growth in both well-irrigated and water-stressed conditions. This effect could be explained by domestication, where increases in biomass due to higher leaf area were observed in several crops (Milla and Matesanz, 2017). Thus, introgression of the wild genotype may represent a backward step in breeding for biomass production. A few ILs had similar growth to the cultivated parent, with only introgressions on chromosomes 6 and 8 standing out in terms of growth for root and stem biomass, respectively. The QTL found on chromosome 6, allowing greater root development under drought conditions, may be of interest, as it has been observed in other species of the genus *Solanum* that genotypes with greater tolerance show greater root development (Manoj and Uday, 2007, Pino et al., 2013). On the other hand, the QTL present on chromosome 8, related to increased stem biomass, was studied in relation to larger xylem size, allowing greater leaf development (Tapia et al., 2015).

The presence of candidate genes for drought tolerance in the ILs evaluated was reported by Gramazio et al. (2017), with 2, 6, 7 and 11 genes present in chromosomes 3, 4, 7 and 12, respectively. However, the ILs containing these introgressions did not show an increase in biomass production under water stress conditions; nevertheless, they could not be discarded as material of interest for breeding, as the introgressions are very large and the study only evaluated the vegetative developmental phase of the crop. For more efficient improvement through introgression, a finer mapping of the QTLs may be required that does not carry undesirable growth-limiting regions. Although the introgressions did not have a positive effect on growth, a QTL related to the maintenance of leaf water content (LWC) during drought (lwc12%) was identified on chromosome 12, which has been considered as a criterion for the selection of drought-tolerant genotypes by other researchers (Rad et al., 2013; Wasaya et al., 2021).

Another trait evaluated was water use efficiency (WUE), which is affected by the effects of climate change, making it crucial to improve this trait through crop selection and cultural practices (Hatfield and Dold, 2019). In the plants evaluated, WUE was positively correlated with growth parameters under both conditions, and a QTL (wue1%) was identified on chromosome 1 that allows for higher WUE when irrigation is reduced. However, it has been shown that higher WUE does not always result in increased yield (Blum, 2009), so further experiments are needed to determine whether this QTL contributes to increased yield.

Water stress in plants can cause a decrease in photosynthetic pigments such as chlorophyll and carotenoids, leading to reduced biomass production (Hosseini et al., 2021; Ibañez et al., 2021). The effect of drought on chlorophyll content in eggplant is not clear, as it has been reported to both increase (Díaz-Pérez and Eaton, 2015) and decrease (Plazas et al., 2019). In this study, although *S. melongena* was largely unaffected by stress, QTLs with a positive effect on photosynthetic pigments were detected in ILs with introgressions on chromosomes 2 and 8. It was observed that under both well-irrigated and water-stressed conditions, photosynthetic pigments were positively correlated with leaf water content, which is another trait reflecting the good physiological status of the plant and is associated with lower yield loss under water stress conditions (Soltys-Kalina et al., 2016).

Proline content under stressful environmental conditions is considered an indicator of the level of stress experienced by plants (Claussen, 2005). All genotypes evaluated, except the wild parent, significantly increased their proline levels in response to stress, as indicated by the PCA, which showed a strong association between higher proline levels and genotypes exposed to water stress conditions. Introgression on chromosome 8 resulted in an even greater increase in proline, which may be associated with drought tolerance due to the high biomass of this genotype. There is evidence that proline content may play a role in the plant’s response to osmoregulation, relative oxygen species (ROS) scavenging, and the regulation of antioxidant metabolites and enzymes during drought stress (Per et al., 2017). On the other hand, the failure to increase proline levels in the wild parent may be due to the adoption of stress avoidance as a drought tolerance mechanism due to slower plant growth and lower evapotranspiration (Fang and Xiong, 2014).

At the molecular level, drought has a major impact by increasing the production of ROS, resulting in oxidative stress in plants (Hasanuzzaman et al., 2013). Malondialdehyde, a compound associated with lipid oxidation, is often used as a marker to assess the level of oxidative stress (Alché, 2019). In the genotypes evaluated, an increase in malondialdehyde levels was observed in stressed plants, suggesting that reduced irrigation led to oxidative stress. Tani et al. (2018) found similar results when studying two eggplant genotypes, which showed increased malondialdehyde and hydrogen peroxide contents under drought conditions. However, in our experiment, the increase in malondialdehyde was only significant in the cultivated parental, so the introgression of the wild parental may have contributed to a higher level of tolerance to oxidative stress, of which MDA is an indicator (Alché, 2019).

In response to oxidative stress, plants can activate enzymatic and non-enzymatic antioxidant systems to eliminate ROS (Demidchik, 2015). In the case of eggplant under water stress, an increase in the activity of enzymes such as superoxide dismutase, catalase, and glutathione reductase was observed (Amiri Rodan et al., 2020). Additionally, phenolic compounds are non-enzymatic antioxidants (Gill and Tuteja, 2010). Total phenolic compounds and flavonoid contents were analysed in this study, but no significant effect was found due to the treatments. Despite this, a positive correlation was observed between these two traits and stem elongation and stem weight, which may suggest that genotypes with higher content possess an antioxidant effect.

## 5. Conclusions

The reduced irrigation regime provided an opportunity to evaluate the impact of water stress on eggplant lines with genetic contributions from the wild relative *S. incanum*, in terms of growth and biochemistry. In general, introgressions of *S. incanum* into the *S. melongena* genetic background result in less growth, and this reduction is, in most cases, counterproductive for water stress tolerance. However, QTLs favouring production under drought, such as increased stem and root biomass, increased leaf tissue hydration, or increased chlorophyll content, were detected. These results are a first step in the analysis of *S. incanum* introgressions, allowing us to identify genomic regions of interest for the genetic improvement of eggplant, particularly for tolerance to drought. The next steps would be evaluating the entire productive cycle of the most interesting ILs to analyse the contribution of introgressions to yield improvement under drought, which is the most important in this type of crop, as well as the fine mapping of the most interesting QTLs to reduce linkage drag and identify candidate genes. In the coming years, new pure ILs carrying the S. incanum introgressions will be developed to increase the altogether donor genome coverage, which could hide unexpected breeding traits such as water stress tolerance.

